# The human mitochondrial 12S rRNA m^4^C methyltransferase METTL15 is required for proper mitochondrial function

**DOI:** 10.1101/809756

**Authors:** Hao Chen, Zhennan Shi, Jiaojiao Guo, Kao-jung Chang, Qianqian Chen, Conghui Yao, Marcia C. Haigis, Yang Shi

## Abstract

Mitochondrial DNA (mtDNA) gene expression is coordinately regulated pre- and post-transcriptionally, and its perturbation can lead to human pathologies. Mitochondrial ribosomal RNAs (mt-rRNAs) undergo a series of nucleotide modifications following release from polycistronic mitochondrial RNA (mtRNA) precursors, which is essential for mitochondrial ribosomal biogenesis. Cytosine N^4^ methylation (m^4^C) at position 839 of the 12S small subunit (SSU) mt-rRNA was identified decades ago, however, its biogenesis and function have not been elucidated in details. Here we demonstrate that human Methyltransferase Like 15 (METTL15) is responsible for 12S mt-rRNA methylation at C839 (m^4^C839) both *in vivo* and *in vitro*. We tracked the evolutionary history of RNA m^4^C methyltransferases and revealed the difference in substrates preference between METTL15 and its bacterial ortholog rsmH. Additionally, unlike the very modest impact on ribosome upon loss of m^4^C methylation in bacterial SSU rRNA, we found that depletion of METTL15 specifically causes severe defects in mitochondrial ribosome assembly, which leads to an impaired translation of mitochondrial protein-coding genes and a decreased mitochondrial respiration capacity. Our findings point to a co-evolution of methylatransferase specificities and modification patterns in rRNA with differential impact on prokaryotic ribosome versus eukaryotic mitochondrial ribosome.

## INTRODUCTION

Mitochondrial gene expression requires a series of inter-connected processes encompassing mtDNA replication and repair, mitochondrial RNA (mtRNA) transcription, maturation and mitoribosome assembly **(1, 2)**. The mt-RNAs, especially rRNAs and tRNAs, are subjected to extensive enzyme-mdiated modifications, which play key roles in RNA stability, RNA structure, and mitochondrial ribosome assembly (**3, 4**). Some of these modifications are deposited co-transcriptionally or immediately after transcription, while others occur when the rRNA is assembled into the pre-ribosomal particle **(3-5)**.

Prokaryotic and eukaryotic cytoplasmic rRNAs contain more than 30 and 200 modified sites, respectively, but only around 10 modifications are found in the mitochondrial rRNAs (**4, 5**). These modifications are located at the functionally important regions of mitoribosome, such as the decoding center (DC) of the small subunit (SSU), suggesting that these modifications might be retained due to their essential roles **(5, 6)**. The best characterized example is the TFB1M-mediated dimethylation on the two highly conserved sites, A936 and A937, at the 3′-end of the mt 12S rRNA, which is necessary for the assembly of the SSU **(7, 8)**. The NOP2/Sun RNA Methyltransferase 4 (NSUN4) forms a complex with MTERF4 to catalyze m^5^C methylation at the position 841 in mt-12S rRNA and to coordinate the mitoribosome assembly (**9, 10**) However, enzymes for m^4^C and m^5^U (uracil) methylation in mammalian mitochondrial rRNAs remain to be identified (**6, 11**).

Mitochondrial diseases may be caused by mutations in mitochondrial DNA (mtDNA) (**12**), but growing evidence suggests that defects in the nuclear genes involved in mitochondrial RNA modifications can also lead to human mitochondrial diseases. For instance, loss of TFB1M results in mitochondrial dysfunction that leads to impaired insulin secretion and diabetes **(13)**. A missense mutation in pseudouridylate synthase 1 (PUS1), which converts uridine to pseudouridine at several mitochondrial tRNA positions, has been reported to be associated with myopathy, lactic acidosis and sideroblastic anaemia (MLASA) **(14)**. Moreover, a defect in the mitochondrial rRNA methyltransferase MRM2 that causes loss of 2’-O-methyl modification at position U1369 in the human mitochondrial 16S rRNA leads to MELAS-like clinical syndrome in patients **(15)**. In addition, an 11p14.1 microdeletion was identified to be highly associated with Attention-Deficit/Hyperreactive Disorder (ADHD), autism, developmental delay, and obesity **(16)**. Intriguingly, the microdeletion region always encompasses a METTL-famlily protein METTL15. More recently, a trans-ancestral meta-analysis of genome-wide association studies uncovered a completely novel single nucleotide polymorphism (SNP) (rs10835310) at METTL15 associated with childhood obesity, (**17**), further implicating the involvement of METTL15 in this human syndrome, although the underlying molecular mechanisms for these associations is still unclear.

In this current study, we demonstrate that human METTL15 protein, encoded by a nuclear gene, is localized in mitochondria and is responsible for methylation of 12S mt-RNA at C839 *in vivo* and *in vitro*. Furthermore, we demonstrate that METTL15-dependent modification of 12S mt-rRNA is necessary for mitoribosome maturation. Our study reveals that methylation of 12S mt-rRNA m^4^C839 by METTL15 is an important epitranscriptome modification, critical for efficient mitochondrial protein synthesis and respiratory function.

## RESULTS

### METTL15 is a mitochondrial protein associated with 12S mt-rRNA

METTL15 is a member of the methytransferase like (METTL) family, characterized by the presence of a binding domain for S-adenosyl methionine, which is a methyl-group donor for methylation reactions **(18, 19**). Through phylogenetic analysis, we found that METTL15 is highly conserved during evolution and is an ortholog of the bacterial methylatranferase, rsmH (**Figure 1A**), which is responsible for the N^4^-methylation of m^4^Cm1402 in 16S rRNA in almost all species of bacteria (**Supplementary Figures S1A**)**(20)**. Given its similarity with rsmH, we wondered whether METTL15 is also a m^4^Cm methyltransferase for rRNA. To address this possibility, we purified SSU rRNA fragments containing C1402 or its equivalent nucleotide from four representative species and measured the levels of m^4^Cm by HPLC-MS/MS. We didn’t detect any meaningful levels of m^4^Cm in the cytoplasmic SSU rRNAs from fruitfly, zebrafish or human, though a large amount of Cm was readily detectable (**Supplementary Figures S1B**), which is consistent with previous reports that the SSU rRNAs of those eukaryotic cells were abundantly modified by 2’-O-methylated cytosine (**Supplementary Figures S1C**)(**21**).

**Figure 1.**
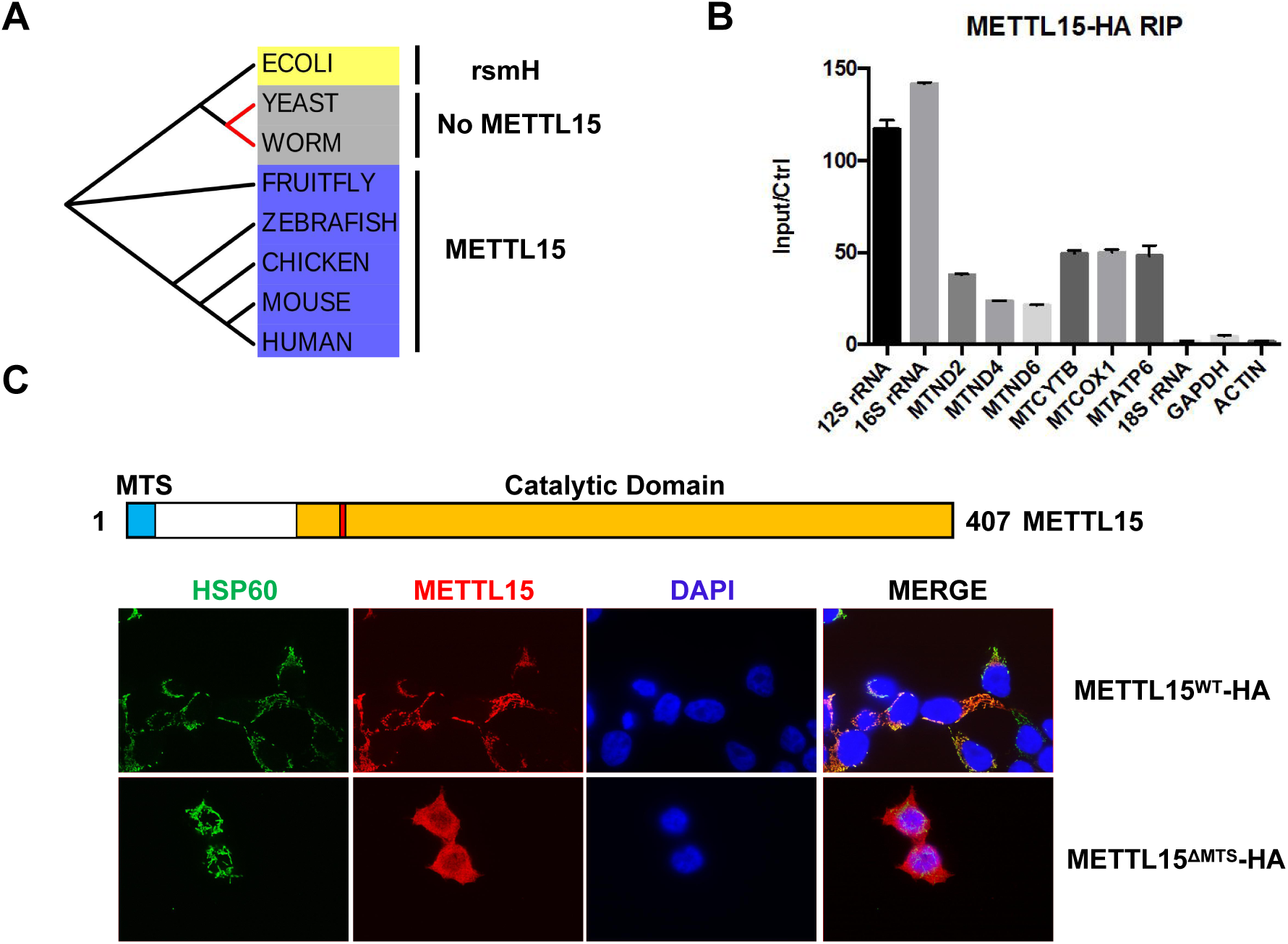
METTL15 localizes in the mitochondria. (A) Phylogenetic analysis demonstrating that METTL15 is likely an evolutionarily conserved m^4^C methyltransferase from fruitfly to human, but absent in worm and yeast; (B) RIP-qPCR analysis of the interactions between METTL15-HA and indicated RNA species. (C) Fluorescence confocal analysis of the subcellular location of wild type, and MTS-deletion METTL5 (Red), HSP60 (Green) was used as a marker for mitochondria.

In eukaryotic cells, mitochondria has its own ribosome translating mitochondrial mRNAs, which prompts us to investigate whether METTL15 is a mitoribosome-specific methyltransferase. Indeed, a considerable amount of mitochondrial genome-encoded RNAs, expecially mt-12S and mt-16S rRNA, but not cytoplasmic RNAs such as 18S rRNAs, are found to be associated with the HA tagged METTL15 in an RNA immunoprecipitation experiment (RIP) (**Figure 1B**). Consistently, immunofluresence experiments showed that METTL15 is exclusively localized in the mitochondria, dependent on its putative mitochondria-targeting signals (MTS) (**Figure 1C**)(**22**), which suggests that METTL15 is a *bona fide* mitochondria protein that interacts with mitoribosome rRNAs.

### METTL15 is responsible for methylation of 12S mt-rRNA m^4^C839 *in vivo*

To unambiguously identify the *in vivo* methylation sites modified by METTL15, we profiled the mitochondrial RNA methylome in wildtype (WT) and METTL15 knockout (KO) cells using RNA bisulfite sequencing (RNA BS-seq), which detects both m^5^C and m^4^C cytosine modifications in RNAs **(23)**. The RNA BS-seq revealed that in the absence of METTL15, the methylation level of mt-12S C839 is dramatically decreased from 58% to near background level (0.9%), which suggests that methylation of mt-12S C839 may be mediated by METTL15 (**Figure 2A and 2B**). To validate the BS-seq results, we designed sequence-specific primers to amplify a 145 nucleotoide (nt) region surrounding C839 from bisulfite-treated RNA samples and employed targeted sequencing (detailed procedure in the Methods) to examine the methylation levels of C839. In agreement with the BS-seq result, methylation of C839 was found to almost completely disappear in the METTL15 KO cells (**Supplementary Figures S2C**). Importantly, the methylation level of m^4^C839 can be fully rescued by wild type METTL15, but not a catalytically compromised mutant METTL15 (GA mutant: _108_GSGG_112_ to _108_ASAA_112_) (**Supplementary Figures S2C)**(**24**), which strongly supports the hypothesis that METTL15 is responsible for m^4^C839 on 12S mt-rRNA *in vivo* and is consistent with a very recent study **(25)**. Interestingly, the neighbouring methylation site, m^5^C841, which is catalyzed by NSUN4 **(9)**, is reduced (but not eliminated) upon METTL15 deletion. In addition, the m^5^C841 reduction could be fully restored by reintroducing wild type METTL15 and partially restored by enzymatically inactive METTL15, suggesting METTL15 might regulate the installation of m^5^C841 by NSUN4 in both enzymatic activity-dependent and independent manner (**Supplementary Figure S2C**).

**Figure 2.**
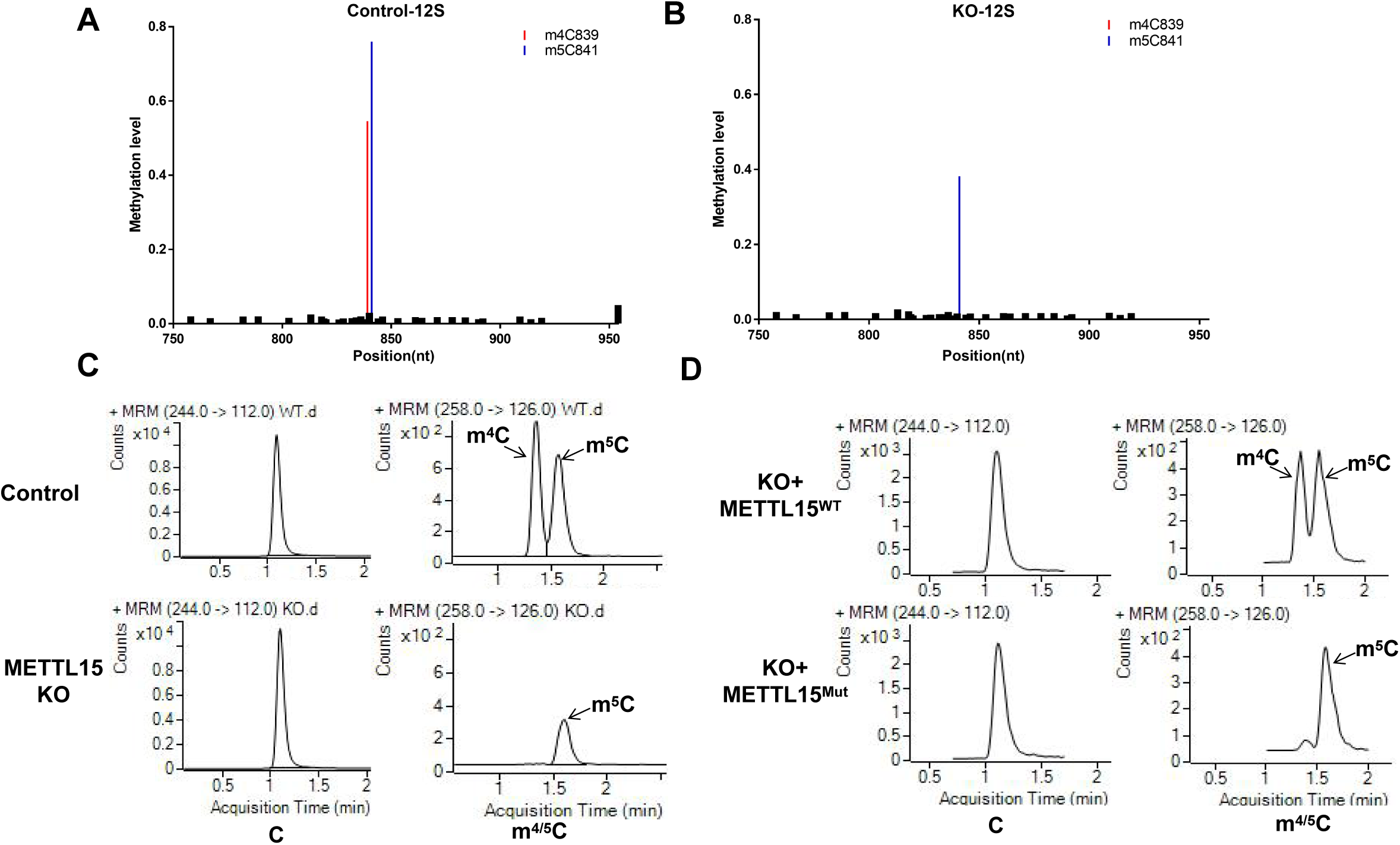
METT15 is the methyltransferase responsible for the m^4^C839 on mitochondrial 12S rRNA *in vivo*. (A) and (B) Relative methylation levels of 12S rRNA determined after sequencing of cDNA obtained from bisulfite treated RNA from control (A) and METTL15 KO (B) cells. (C) LC-MS/MS chromatograms of C, m^4^C, and m^5^C in the corresponding 12S rRNA fragments purified from total RNA. Samples from Ctrl, METTL15 KO cells were analyzed. (D) The m^4^C methylation is readily restored by re-expression of WT METTL15 but not the catalytic mutant.

Given both m^4^C(m) and m^5^C are able to block the C-to-T transition by bisulfite treatment and therefore can’t be distinguished in the BS-seq analysis (**23**), we turned to an optimized LC-MS/MS method to efficiently separate different forms of methyl cytosines in order to define the exact type of methylation in C839 (**Supplementary Figures S2B**). As shown in **Figure 2C**, m^4^C was detected in WT cell lines but reduced to a background level in the METTL15 KO cells, indicating that METTL15 may be a m^4^C methyltransferase for C839. Consistent with the BS-seq data, HPLC-MS/MS analysis also found a modest reduction of m^5^C at C841 due to METTL15 depletion, which again points to a potential crosstalk between C839 and C841 methylation. m^4^C methylation of C839 is mediated by the intrinsic enzymatic activity of METTL15 as reintroduction of wildtype, but not the catalytically inactive METLL15, back into the METTL15 KO cells restored the methylation level of m^4^C839 (**Figures 2D** and **Supplementary Figures S2A/C**). Collectively, these findings demonstrate that METTL15 is likely the enzyme responsible for methylation of mt-12S m^4^C839 *in vivo* (**Supplementary Figures S2D**).

### METTL15 methylates 12S mt-rRNA m^4^C839 *in vitro*

We next asked whether recombinant METTL15 mediates 12S mt-rRNA methylation at C839. C-terminal FLAG-tagged METTL15 was expressed in 293T cells and purified using an ANTI-FLAG M2 Affinity column (**Supplementary Figures S3A**). Recombinant METTL15 was incubated with 12S mt-rRNA oligos (nt 832 to 846) in the presence of d3-SAM (S-(5’-Adenosyl)-L-methionine-d3) as a methyl group donor, and the resulting rRNAs oligos were isolated for LC/MS measurement. As shown in **Figure 3A**, m^4^ methylation of C839 was successfully detected while the catalytically compromised METTL15 failed to mediate C839 methylation.

**Figure 3.**
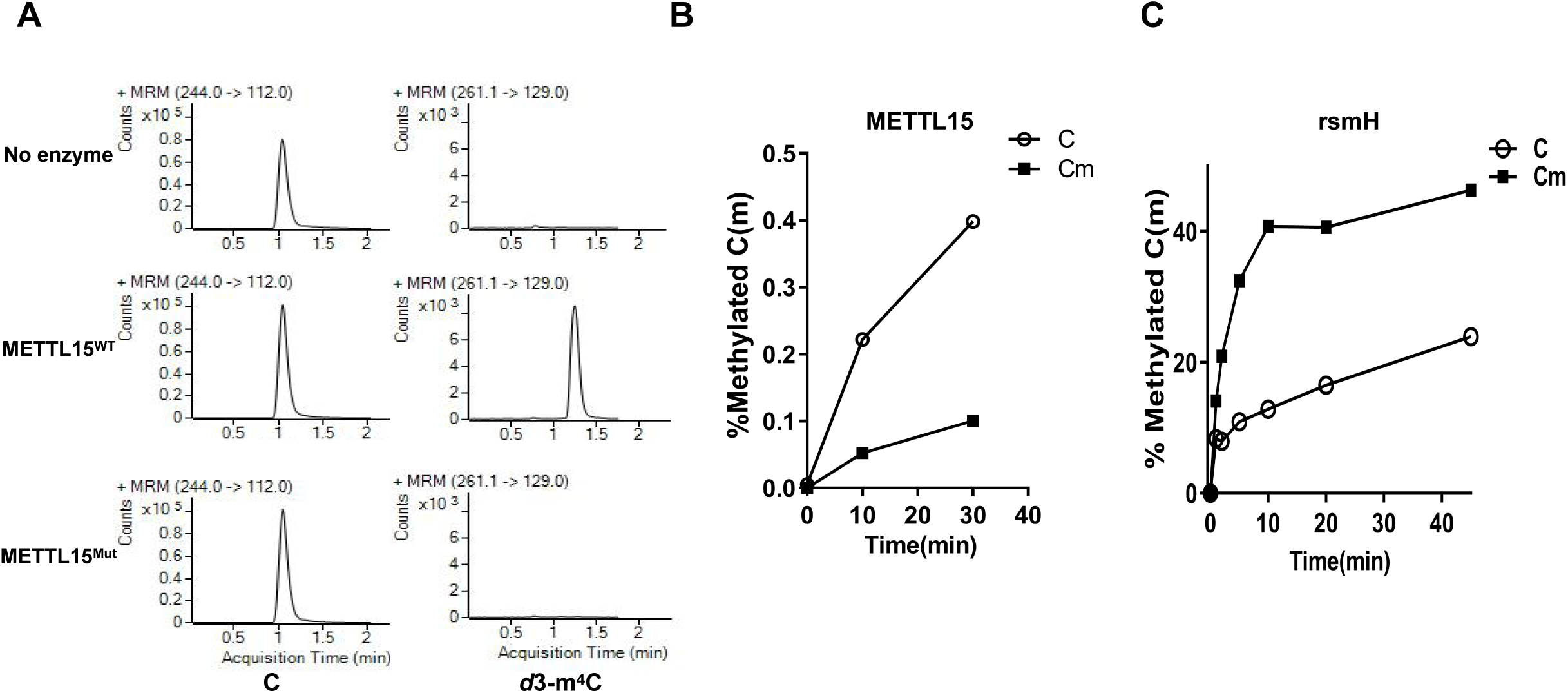
METT15 is the methyltransferase responsible for the m^4^C839 on mitochondrial 12S rRNA *in vitro*. (A) *In vitro* methylation assay indicates that recombinant wild-type, but not catalytic mutant METTL15, is able to deposit a methyl group onto the N^4^-position of C839 in 12S mt-rRNA. (B) Human METTL15 prefers RNA oligos with unmodified C839. (C) Bacterail rsmH methylatransferase shows stronger activtiy toward 2’-O methylated subsrates.

As described before, a main difference between the m^4^C methylation site of bacterial ribosome and human mitoribosome is that cytosine in bacterial ribosome is mainly m^4^Cm while in the human mitoribosome at the equvalent cytosine residue, it’s m^4^C without the 2’-O methylation. This prompted us to determine whether human MEETTL15 (hMETTL15) displays any preference for unmodified cytosine versus 2’-O methylated cytosine. Consistently, unmodified cytosine in the mt-12S rRNA appears to be a better substrate for hMETT15 *in vitro* (C vs Cm in **Figure 3B**). In contrast, rsmH shows higher apparent activity towards Cm compared with C of mt 12S rRNAs (solid line in **Figure 3C, Supplementary Figures S3C)**. Interestingly, we found an aromatic amnio acid (W139) in rsmH, which has been changed into Valine (V210) in the eukaryotic orthologs (METTL15), might potentially mediate the interaction of 2’-O methyl group of Cm with rsmH based on the published rsmH structure (**Supplementary Figures S3B**) (**26**). These results confirmed that human METTL15 is a *bona fide* m^4^C methyltransferase and has a higher activity toward unmodified cytosine compared with 2’-O methylated cytosine *in vitro.*

### Depletion of METTL15 inhibits the function of mitoribosomes

As METTL15 is localized in mitochondria, we first investigated the effect of METTL15 deletion on mtDNA copy number and transcription of mitochondrial genome-encoded genes. We found METTL15 deletion only causes minor changes of mtDNA copy number and transcription (**Figure S4A/B**). Given that the methylation site lies in the critical region of mitoribosome, we asked whether loss of METTL15 affects the function of mitoribosome. We performed the mitochondiral ribosome profiling in a 10%-30% sucrose gradient to determine whether there was any difference in the assembly of mitoribosome. The distribution of SSU and LSU in the sucrose gradient were detected by the presence of 12S and 16S mt-rRNA, respectively. According to the protein complex density, the first peak of 12S rRNA (fraction 8) represents SSU, while the first peak of 16S rRNA (fraction 9) represents LSU, and the co-fractionated peaks (fractions 12 and 13) represent the mature ribosome. The co-fractionation ratio of 12S and 16S mt-rRNAs in METTL15 KO cells was significantly reduced (compared with factions 12-13 in wildtype cells), thus identifying a major defect in mitoribosome assembly. In addition, the ratio of mRNA encoded by mitochondrial genome was also significantly reduced in the 55S mature monosomes (fractions 12-13), indicating compromised translation efficiency, which was consistent with the observed mitoribosome assembly defects (**Figure 4A and 4B**). Western blotting results of two representative mitochondrial protein-coding genes, COX2 and ND6, showed that the levels of the protein products were also reduced significantly. (**Figure 4C**). Importantly, the translational defects of the mitochondria-encoded genes could be rescued by wild type METTL15 but not the catalytic mutant (**Figure S4D**), suggesting the function of METTL15 on mitoribosome is m^4^C849 dependent. These data thus demonstrate that the methylation mediated by METTL15 is critical for mitoribosomes maturation and mt-mRNA translation.

**Figure 4.**
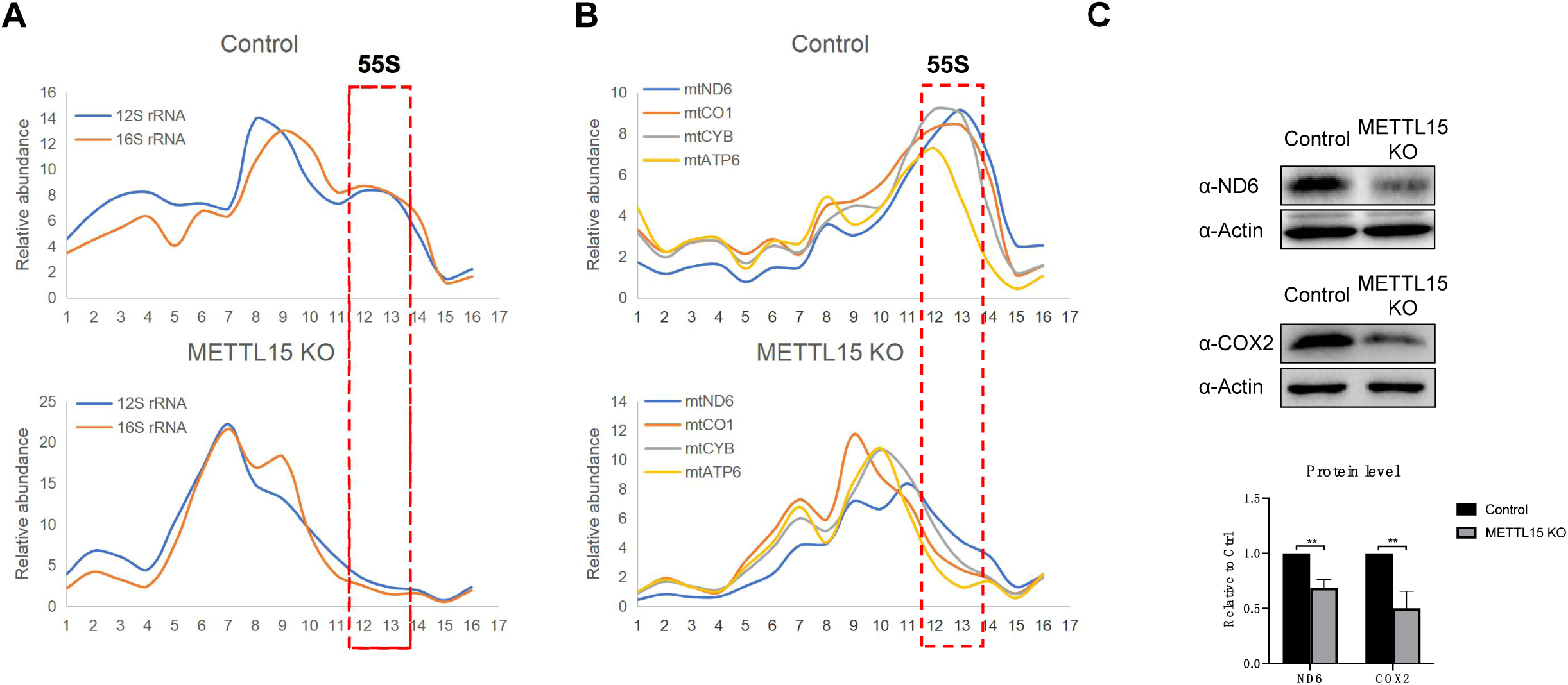
METTL15 is required for proper mitochondrial ribosome assembly and the translation of genes encoded in the mitochondrial genome. (A) The distribution of the mitochondrial ribosome small and large subunits in the indicated sucrose gradient fractions, examined by 12S and 16S rRNA RT-qPCR. (B) The distribution of the mRNAs of the mitochondrial coding genes in the indicated sucrose gradient fractions examined by RT-qPCR. (C) Western blot analyses of ND6 and COX2 protein levels in the control and METTL15 KO cells. The bar graph represents the quantification results of 2 replicate experiments.

### Proper functions of mitochondria are affected by METTL15 depletion

The most prominent role of mitochondria is to produce adenosine triphosphate (ATP)-through respiration, and to regulate cellular metabolism. Most of the ATP synthesized during glucose metabolism is produced in the mitochondria through oxidative phosphorylation (OxPhos) powered by the Electron Transport Chain (ETC) complex, which consists essentially of about 70 nuclear-encoded proteins and 13 mtDNA-encoded proteins translated by mitoribosome in mitochondria. To determine the impact of loss of METTL15 on respiratory activity, we measured the respiratory activity of METTL15 KO cells using a Seahorse XF96 analyzer. The oxygen consumption rate (OCR) of the METTL15 KO cells was substantially lower than that of WT cells, and this effect is dependent on the enzymatic activity, indicating that METTL15 mediated m^4^C839 on 12S mt-rRNA is required for proper oxidative phosphorylation function. (**Figure 5A/B**). After two days in culture, the medium of METTL15 KO cells turned more yellow indicating a lower pH and more lactate secretion, although the cell numbers are comparable between WT and KO, (**Figure S5A**). Consistently, extracellular acidification rate (ECAR), which approximates glycolytic activity, was significantly up-regulated in the METTL15 KO cells, likely to compensate for dysfunction of the mitochondria (**Figure 5C**). Furthermore, metabolites profiling shows a decline of citrate and alpha-ketoglutarate, the intermediators of the TCA cycle, which is closely coupled with OxPhos to generate ATP. It is also known that an essential function of respiration in proliferating cells is to support aspartate biosynthesis **(27, 28)**. The decline of the aspartate level in METTL15 KO cell is consistent with the compromised respiration function. At the same time, the upregulated level of lactate suggested that cells use more anaerobic glycolysis to compensate for impaired mitochondrial function (**Figure 5D/S5A**). These results indicate that METTL15 is important for maintaining mitochondrial function and cellular metabolic homeostasis.

**Figure 5.**
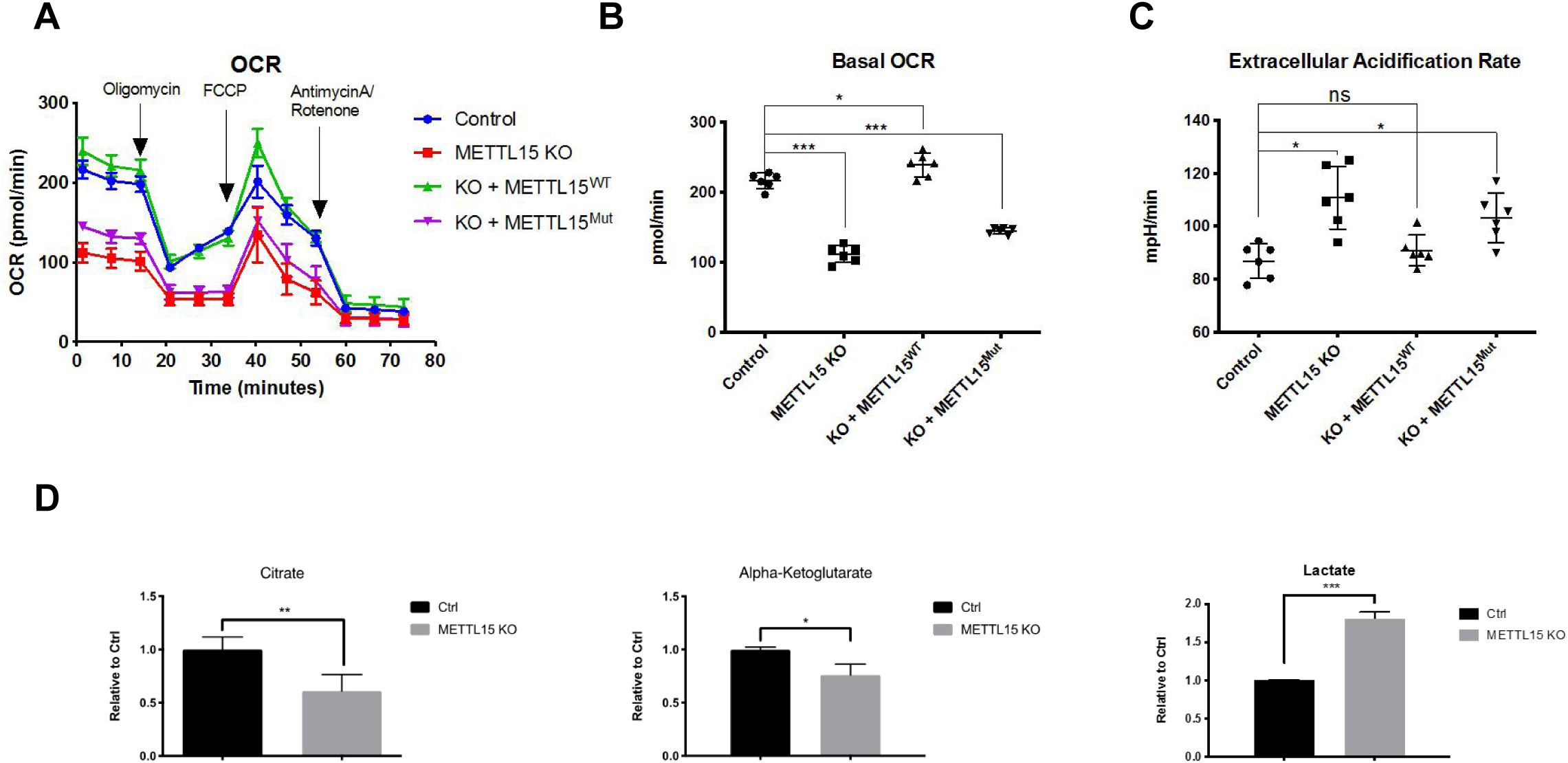
METTL15 deletion facilitates the transformation from aerobic glycolysis to anaerobic glycolysis. (A) Oxygen consumption rate (OCR) of the control, METTL15 KO cells and the METTL15 KO cells with the wild type or catalytic mutation of METTL15-Flag-HA rescuing constructs measured by Seahorse XF96 machine. (B) Quantification of the basal OCR for the indicated cells. (C) Quantification of extracellular acidification rate (ECAR) for the indicated cells at the basal condition. (D) Cellular metabolite concentration determined by liquid chromatography-tandem mass spectrometry (LC-MS/MS) in the control and METTL15 KO cells. All data are represented as mean ± SD from four biological replicates. * p < 0.05; ** p < 0.01; *** p <0.001, T test.

## DISCUSSION

Here we describe the identification of METTL15 as the methyltransferase that generates m^4^C839 in human 12S mt-rRNA. The phylogenetic analysis indicates that METTL15 orthologs exist in most eukaryotes, which implies the importance of this methyltransferase for proper functions of mitoribosome. Consistent with this hypothesis, mitochondrial translation is inhibited and oxidative phosphoryaltion is remarkably compromised in METTL15 KO cells, identifying an important function of METTL15 in regulating mitochondria functions by methylating 12S mt-rRNA.

### Crosstalk between m^4^C839 and m^5^C841 on 12S mt-rRNA

Ribosomal maturation involves multiple steps of subunit assembly and incorporation of chemical modifications into the rRNA **(3)**. The assembly of protein components and rRNAs has been well characterized through high-resolution Cryo-EM, while the process of modification deposition is still largely unclear **(29)**. For the bacterial 16S rRNA, quantitative analysis of rRNA modifications finds that the modification events seem to occur in a 5′-to-3′ sequential order: from 5′ body domain, the 3′ head domain, to 3′ minor domain **(30)**. In this current study, we found that m^4^C839 methylation appears to precede m^5^C841 and important for the nearby m^5^C841 methylation, suggesting crosstalk between modifications of the two nearby residues. Furthermore, the enzymatically inactive METTL15 can partially restore the m^5^C841 methylation decreased in METTL15-null cells, raising the question of whether the crosstalk is mediated by physical interactions between METTL15 and the m^5^C methyltransferase, NSUN4. Undoubtedly, the investigation of how modifications of mt-rRNA are coordinately deposited in an orderly manner will significantly increase our understanding of mitoribosome maturation **(31)**.

### Co-evolution of rRNA methyltransferases and rRNA fuctions

Unlike the universally conserved rRNA modifications (such as m^6,6^A) (**5**), the N^4^-methylation of SSU rRNA is only maintained in prokaryotes and mitochondria of eukaryotic cells. In bacteria, rsmI and rsmH (which is the bacterial METTL15 ortholog) install 2’-O methylation and N^4^-methylation, respectively, on the equivalent cytosine to generate m^4^Cm1402 of bacterial 16S rRNAs **(20)**. There are two notable differences between methylation of human mt-rRNAs versus bacterial rRNA. First, at the molecular level, our *in vitro* enzymatic assays showed that rsmH prefers Cm as a substrate and m^4^C methylation at C1402 in bacteria may occur subsequent to the 2’-O methylation. Interestingly, the aromatic pocket in the rsmH enzymatic domain appears to have degenerated in its eukaryotic ortholog METTL15 proteins during evolution, and this might be coupled with the loss of rsmI in eukaryotic organisms. Second, the m^4^C methylation is essential for the mitoribosome assembly and maturation, while in contrast, m^4^Cm loss in bacteria only generates a rather modest phenotype (**20**). These findings suggest that m^4^C appears to have gained importance in regulating mitochrondrial functions during evolution.

### METTL15 and human diseases

Previous studies demonstrate that mitochondrial dysfunctions in mature adipocytes cause defects in fatty acid oxidation, secretion of adipokines, and dysregulation of glucose homeostasis **(32)**. Importantly, a reduction in the oxidative capacity of brown adipocytes results in impaired thermogenesis, and has been linked to diet-induced obesity **(33)**. In this context, it’s important to note that microdeletion and SNPs in METTL15 gene are highly associated with obesity **(16, 17)**. Therefore, we speculate that if altered METTL15 activity impacst obesity onset, it’s likely to be due to the ability of METTL5 to regulate mitochondrial functions by methylating 12S mt-rRNA.

In summary, we identify METTL15 as a m^4^C methyltransferase that specifically mediates m^4^C methylation of 12S mt-rRNA at residue C839. Interestingly, our HPLC-MS/MS and enzymological analyses reveal a methylation pattern shift during evolution, which is likely a consequence of degeneration of an ancestral aromatic pocket present only in bacterial METTL15 orthologs, which recognizes 2’-O-methyl cytosine. This pocket is absent in eukaryotic METTL15, which is probably why human METTL15 prefers C839 but not Cm839. Importantly, METTL15 depletion affects mitoribosome assembly, inhibits translation of mitochondria encoded proteins, and compromises the oxidative phosphorylation, which underlies the importance of METTL15 in maintaining mitochondrial functions. Future experiments will determine whether METTL15 plays a role in human diseases such as obesity through its ability to mediate methylation of mitochondrial rRNA.

## METERIALS AND METHODS

### Constructs and Antibodies

For the expression of METTL15 protein, METTL15 ORF cDNA was cloned into pHAGE or pET28a expression vector (Invitrogen). METTL15 and rsmH mutants were generated using the QuikChange Site-Directed Mutagenesis Kit (Stratagene) according to the manufacturer’s protocol.

Anti-FLAG (M2) beads and antibody (F1804) were purchased from Sigma. Total OXPHOS Human WB Antibody Cocktail (ab110411) and anti-ND6 (ab81212) antibodies were purchased from Abcam. Anti-COX2 (55070-1-AP)antibody was purchased from Proteintech.

### Immunofluorescence

Cells were seeded in 24-well plates at a density of 20,000 cells/well one day before IF examination. Through standard fixation, permeabilization as well as antibody incubation procedures, confocal imaging was taken by Yokogawa spinning disk confocal on an inverted Nikon Ti fluorescence microscope and then processed by Image J.

### Generation of METTL15 knockout cell lines

The CRISPR-Cas9 targeting system was utilized as previously described **(34)**. The guide RNA sequence is *METTL15* KO: 5’-TTGAGATCTGTGTAACTCCT-3’, targeting exon 3 of the NCBI *METTL15* reference sequence NM_001113528.

### RNA immunoprecipitation (RIP)

A total of 5 million METTL15-HA stable cells were crosslinked with 1% formaldehyde and then harvested and lyzed in 3 volumes of lysis buffer (50 mM Tris-Cl [pH 7.5], 300 mM NaCl, 0.5 mM DTT, 0.25 % NP-40 with protease inhibitor) on ice for 15 min. After centrifugation at 14,000 rpm for 15 min at 4 °C, the supernatant was used for IP. For each RIP reaction, 5 million cells and 10 μl HA beads were used. After washing, the precipitated RNA samples were extracted by TRIzol directly and reverse transcribed, followed by qPCR detection.

### *In vitro* RNA methylation assay

Recombinant rsmH wildtype and mutant proteins were expressed in the Rosetta (DE3) bacterial cells, which were incubated at 37 °C until OD600 reached ∼ 0.6–1 and then cooled down to 16 °C. IPTG was added to 0.2 mM final concentration, and cells were further incubated at 16 °C for 16 h. Cell pellets were lysed in a buffer containing 300 mM NaCl, 25 mM Tris pH 7.5, 10% Glycerol, 0.5% NP-40. Total lysate was incubated with HisPur Ni-NTA Resin at 4 degree for 5 hours. His-rsmH protein was eluted with elution buffer (20 mM Tris pH7.5, 150 mM NaCl, 200mM Imidazole) in 0.5 ml aliquots until color change was no longer observed (Bradford assay).

### Sucrose gradient sedimentation

Sucrose gradient sedimentation was performed as previously described with minor modifications **(35)**. For each sample, around 1 mg mitochondria protein lysate was used, and the lysates were loaded on a 10 ml 10 % – 30 % uncontinuous sucrose gradient (50 mM Tris-Cl, 100 mM KCl, 10 mM MgCl_2_) and centrifuged at 32,000 rpm for 130 mins in a Beckman SW60-Ti rotor. A total of 16 fractions were collected from the top and used for RT-qPCR analyses.

### Seahorse assay

An XF96 extracellular flux analyzer (seahorse bioscience) was used to determine the oxygen consumption rate(OCR) between WT and METTL15 KO cells, which were seeded at 15,000 cells per well. The concentrations of oligomycin, FCCP, rotenone and antimycin A are 1 μM, 1.5 μM, 0.5 μM and 0.5 μM, respectively.

### Isolation of a defined rRNA fragment

To isolate the corresponding fragments of 12S rRNA containing known modified residues, we refer to the method as previously described **(36)** with minor modifications. Briefly, we used 200 pmol of biotin-tagged synthetic oligodeoxynucleotide probe and 100 μg of total RNA for one experiment and the sequences of the probes are listed in the supplementary table. After annealing and digestion with mung bean nuclease (NEB) and RNase A, the duplex of the probe and corresponding RNA fragment is purified with streptavidin C1.

### Processing of RNA samples for mass spectrometry

The RNA sample is digested with 0.5U of Nuclease P1 (Sigma) in 80 μl NP1 buffer at 42°C for 2 hours. To dephosphorylate the single nucleotides, 1U of CIP (NEB) is added and incubated for an hour at 37°C. The samples are filtered with Millex-GV 0.22 μm filters before loaded onto the column of mass spec machine.5 μl of solution was loaded into LC–MS/ MS (Agilent6410 QQQ triple-quadrupole mass spectrometer). Nucleosides were quantified by using retention time and nucleoside to base ion mass transitions of 244 to 112 (C), 258 to 126 (m^4^C and m^5^C), 258 to 112 (Cm) and 272 to 126 (m^4^Cm).

### Bisulfite mapping of mC residues in mitochondrial RNA

For purified mitochondria, mitochondrial RNA for bisulfite treatment was isolated with TRIzol and treated with TURBO DNase (Ambion) to remove mitochondrial DNA. Bisulfite treatment was performed with the EZ RNA methylation kit (Zymo research). Half of the treated RNA was used for bisulfite RNA-seq library preparation with NEBNext Ultra II Directional RNA Library Prep Kit (NEB), according to the manufacturers instructions. For the targeted bisulfite analysis, the bisulfite-treated RNA was directly converted to cDNA using PrimeScript RT Reagent Kit(Takara Bio, Inc.), followed by PCR amplification using primers specific for the region surrounding C839 of 12S mt-rRNA. The PCR products were separated from unincorporated primers using low melting agarose and submitted for Amplicon-EZ sequencing (Genewiz).

BS RNA-seq was carried out on Illumina NextSeq platform with single-end 75 bp read length. Raw reads were stripped of adaptor sequences and removed of low quality bases using Cutadapt. The processed reads were aligned to human mitochondrial genome with meRanT align(meRanTK, version 1.2.0) and the methylation ratio was calculated by meRanCall (meRanTK, version 1.2.0)(**37**).

## ACKNOWLEDGEMENTS

We thank Alison Ringel and Kiran Kurmi for assistance with setting up the Seahorse experiments, Phillip A. Dumesic, Marwan Shinawi and Alberto Fernández Jaén for their insightful advice, and Noa Liberman-Isakov in Eric Greer’s lab for providing the N^4^-Methyl-CTP standard. We also would like to thank Santa Cruz Biotechnology, Inc. and Abclonal, Inc. for providing antibodies.

## Authors contributions

Y.S. and H.C. conceived and designed the project. H.C., Z.S., K.C., C.Y. and Q.C. carried out the major experiments. H.C. and J.G. performed the bioinformatic analysis. M.H. supervised the analysis concerning the functions of METTL15 in mitochondria. Y.S. supervised the project throughout. Y.S., H.C. and Z.S. co-wrote the manuscript and all authors contributed to the manuscript writing.

## Figure Legends

**Figure S1.**
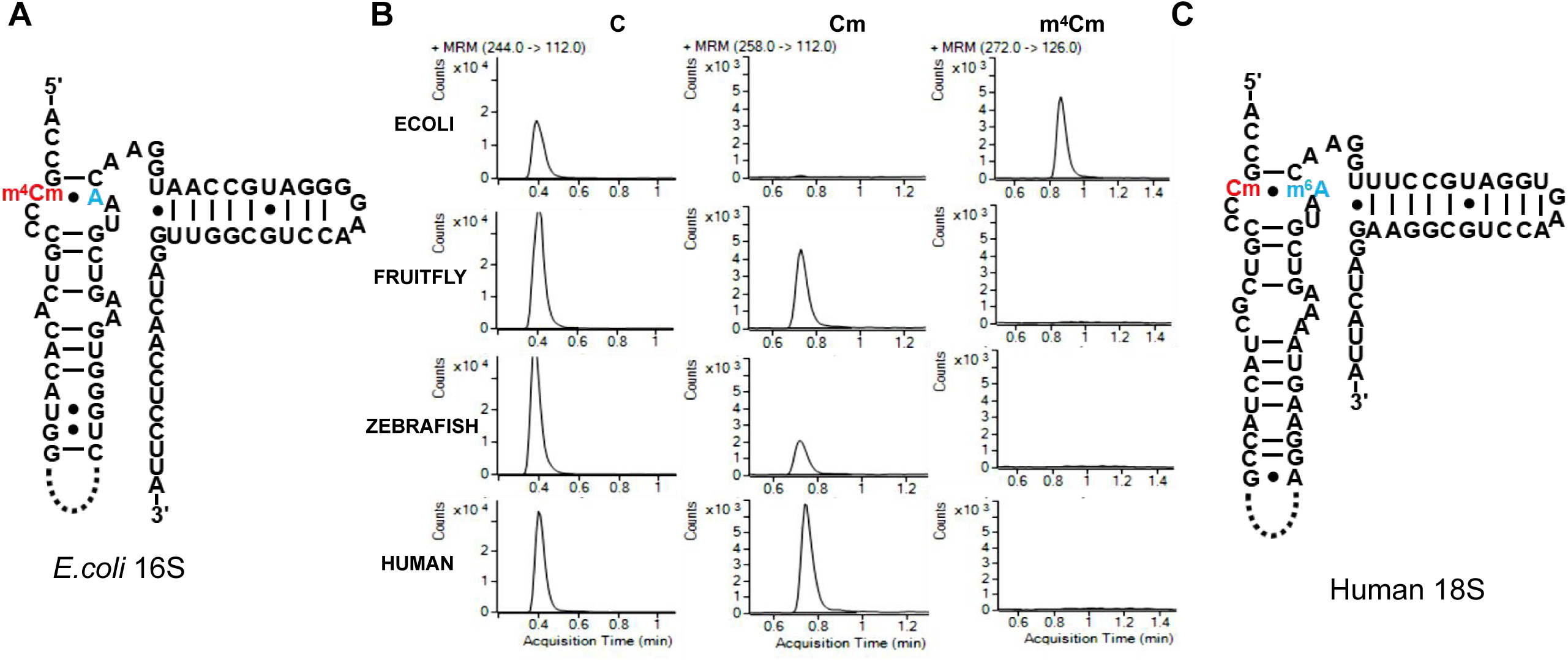
(A) The bacterial m^4^Cm is localized in h44 of SSU rRNA. (B) LC–MS/MS quantification of m^4^Cm in SSU rRNA demonstrating that considerable amount m^4^Cm on small subunit rRNA can be detected only in bacteria. (C) The human SSU rRNA bears Cm and m^6^A but not m^4^Cm in h44 of SSU rRNA.

**Figure S2.**
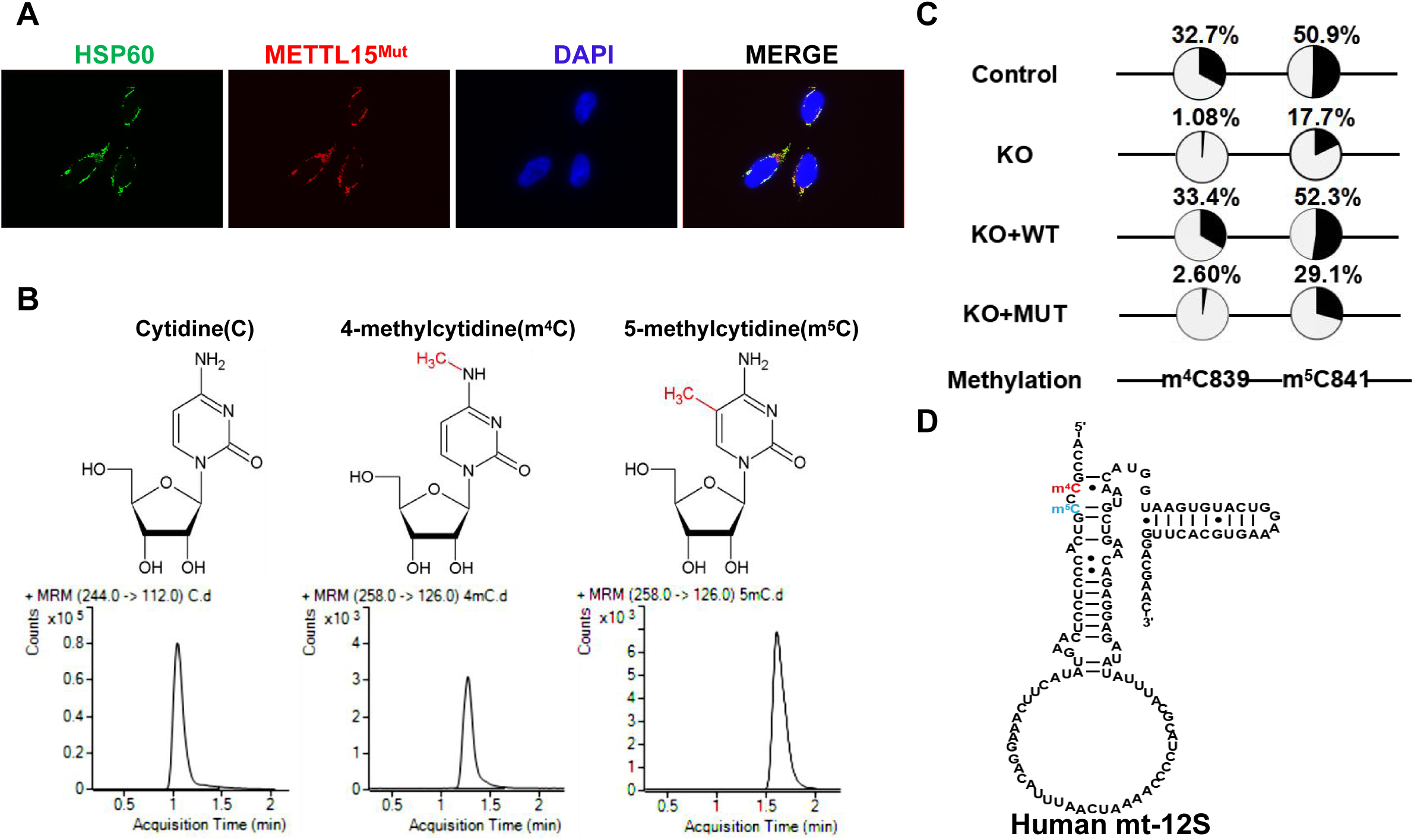
(A) The mutation in the catalytic activity of METTL15 doesn’t affect its mitochondrial localization. (B) LC-MS/MS chromatograms of C, m^4^C, and m^5^C standard. (C) Targeted BSP sequencing shows that METTL15 KO induces a dramatic reduction of m^4^C839 and m^5^C841 methylation, which is readily rescued by wild-type METTL15. In contrast, m^4^C839 can not be restored by the catalytic mutant of METTL15 at all while m^5^C841is partially restored. (D) The secondary structure of human SSU mt-rRNA 3’ terminus and the localization of m^4^C and m^5^C.

**Figure S3.**
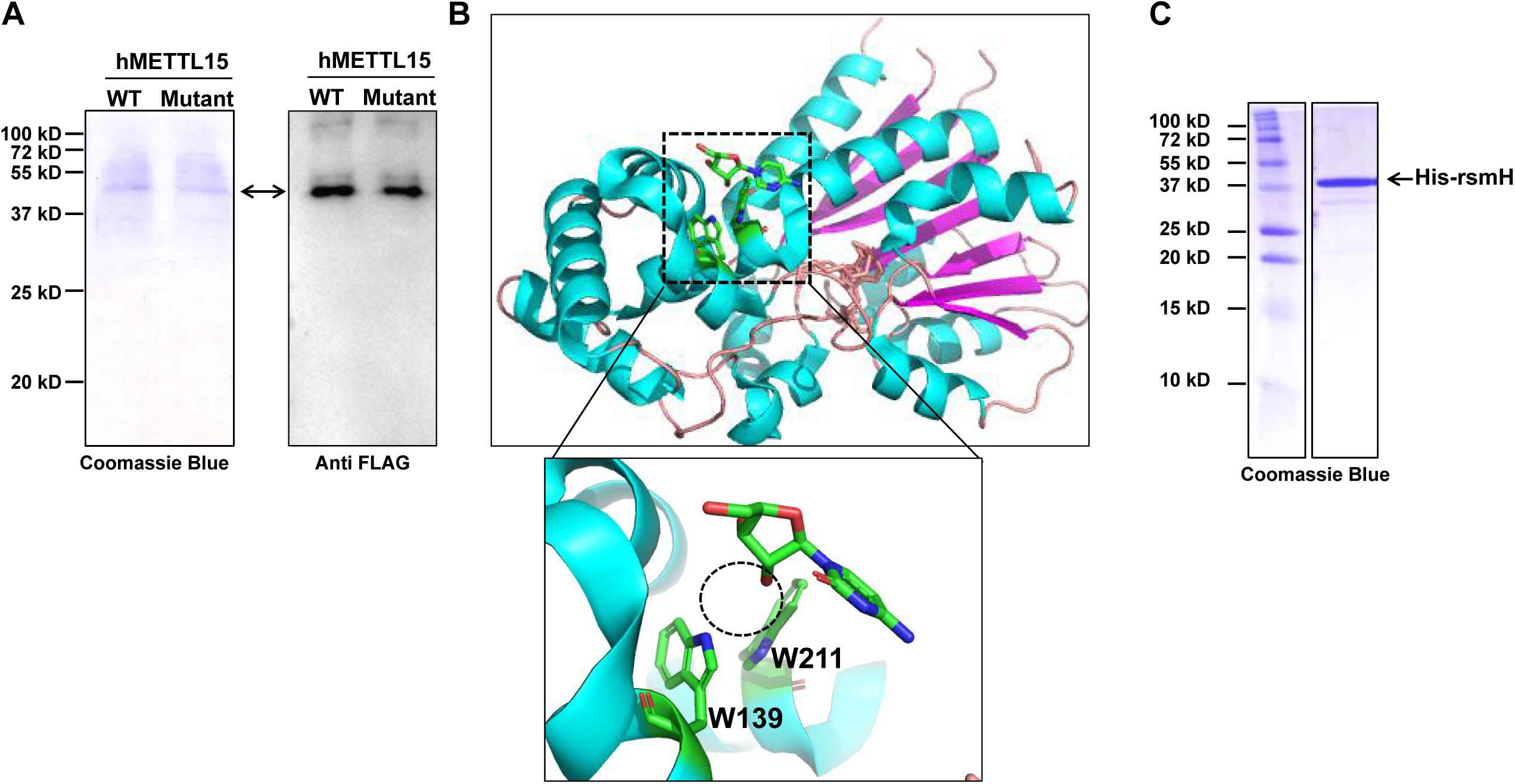
(A) Validation of recommbinant METTL15 purification by Coomassie Blue staining(left) and Western blotting (right). (B) Analysis of the crystal structure of rsmH shows that rsmH has a aromatic pocket that might be involved in recognizing 2’-O methylation of 2’-O-methylcytidine. Note: cytidine was used in the original co-crystal structure. The dash circle indicates the predicted position of 2’-O methyl group when rsmH uses 2’-O-methylcytidine as substrate. (C) Purification and Coomassie Blue staining of purified recommbinant bacterail rsmH.

**Figure S4.**
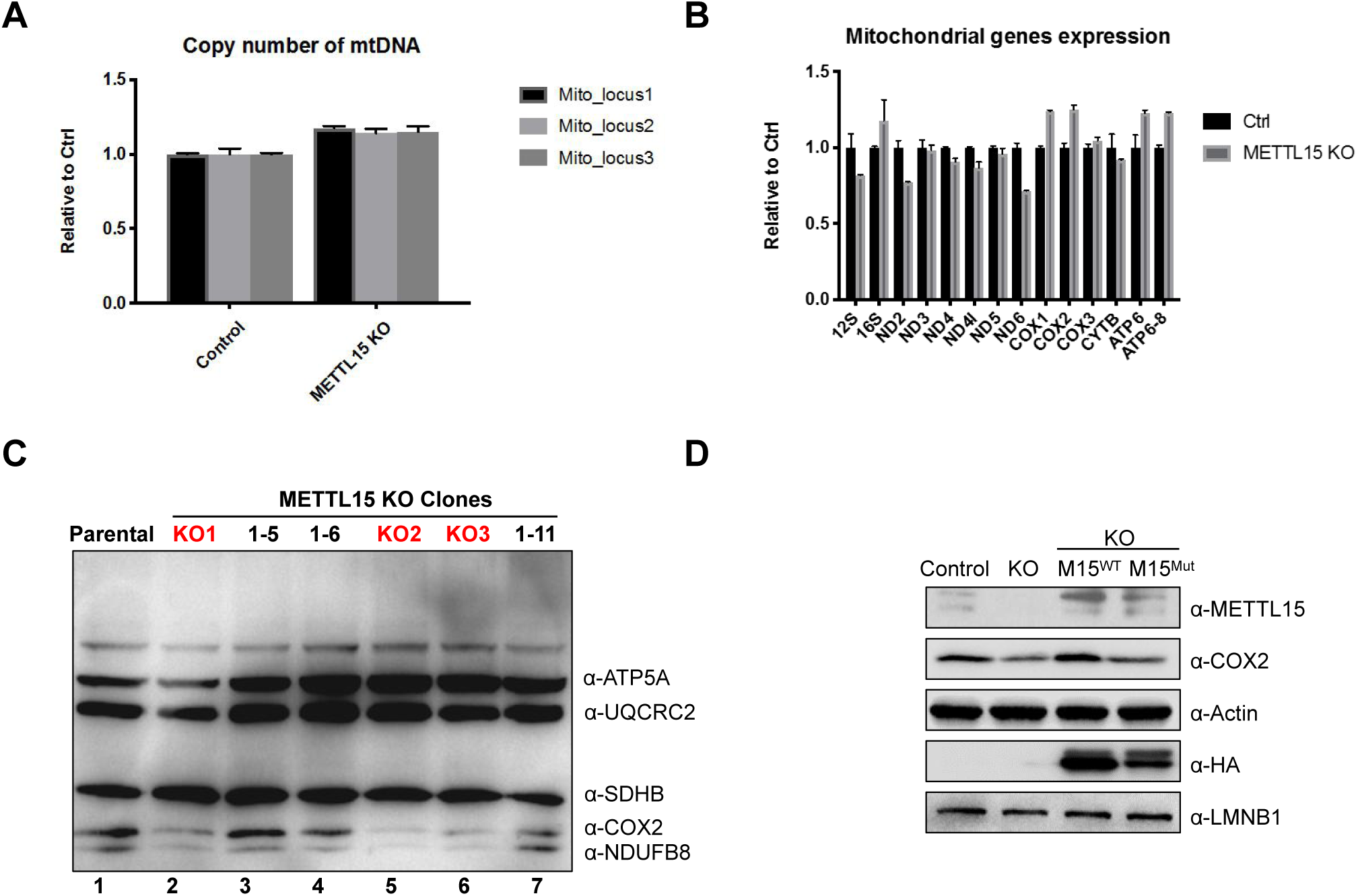
(A) Comparison of mitochondrial genome (mtDNA) copy number between the Control and METTL15 KO cells, measured by RT-qPCR. (B) Expression analysis by RT-qPCR of mitochondrial coding genes in the control and METTL15 KO cells. (C) The protein level of COX2 is downregulated in successfully edited cell lines (red fonts) but not changed in parental or unedited cells (black fonts). (D) Western blot analysis of the indicated proteins in the control, METTL15 KO, and METTL15 KO cells rescued with the wildtype or catalytic mutant of METTL15-Flag-HA, respectively. LMNB1 and Actin were used as controls.

**Figure S5.**
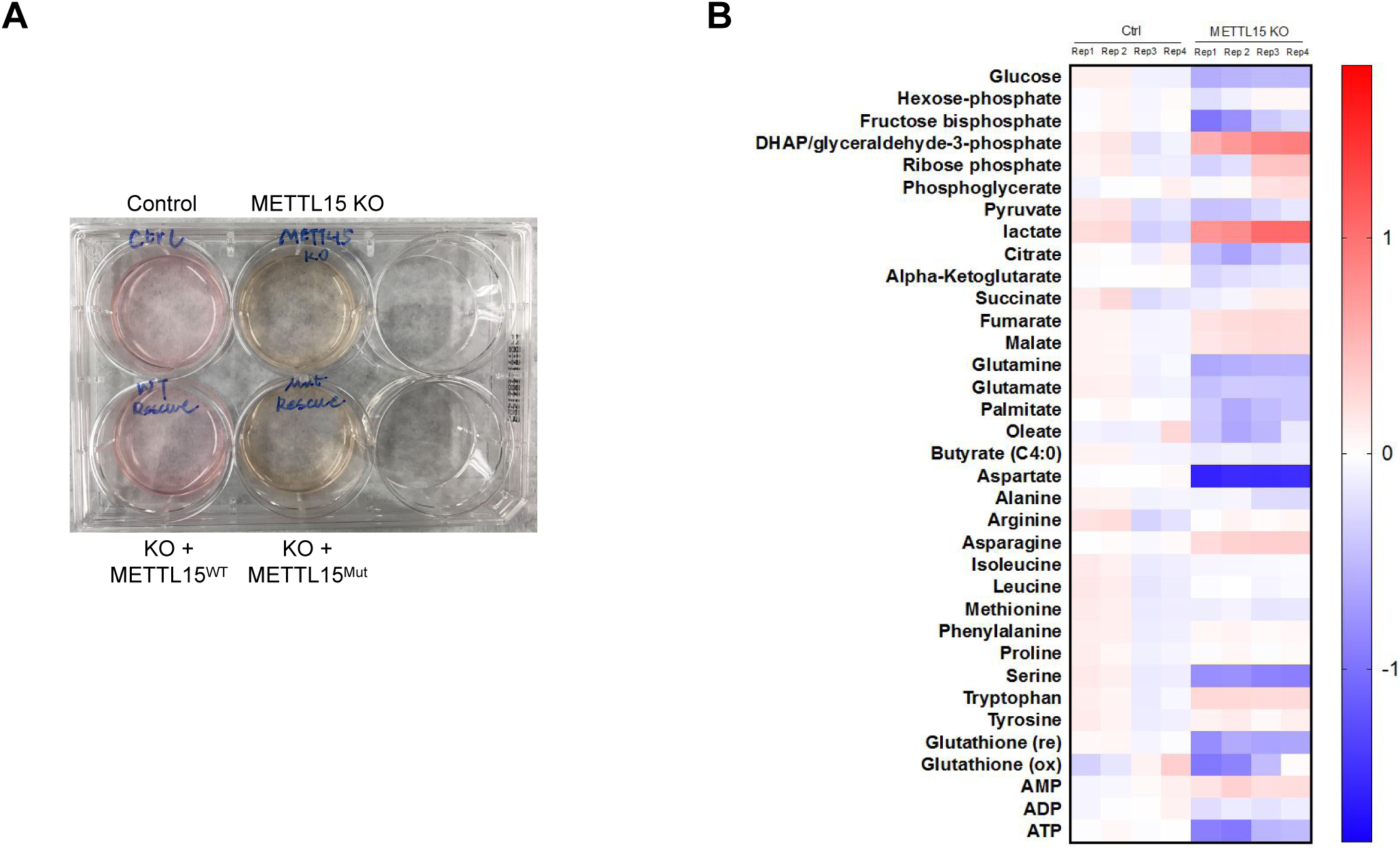
(A) Images showing different medium colors of METTL15 KO cells rescued with either wildtype or catalytic mutant of METTL15, two days after the same numbers of cells were seeded. (B) Heatmap of the metabolites abundance change for control and METTL15 KO cells. Levels of various metabolites were acquired by LC-MS/MS.

## RERFENCE

1. Costanzo MC & Fox TD (1990) Control of mitochondrial gene expression in Saccharomyces cerevisiae. Annual review of genetics 24:91–113.

2. Taanman JW (1999) The mitochondrial genome: structure, transcription, translation and replication. Biochimica et biophysica acta 1410(2):103–123.

3. Pearce SF, et al. (2017) Regulation of Mammalian Mitochondrial Gene Expression: Recent Advances. Trends in biochemical sciences 42(8):625–639.

4. Bohnsack MT & Sloan KE (2018) The mitochondrial epitranscriptome: the roles of RNA modifications in mitochondrial translation and human disease. Cellular and molecular life sciences: CMLS 75(2):241–260.

5. Sergiev PV, Aleksashin NA, Chugunova AA, Polikanov YS, & Dontsova OA (2018) Structural and evolutionary insights into ribosomal RNA methylation. Nature chemical biology 14(3):226–235.

6. Dubin DT (1974) Methylated nucleotide content of mitochondrial ribosomal RNA from hamster cells. Journal of molecular biology 84(2):257–273.

7. Metodiev MD, et al. (2009) Methylation of 12S rRNA is necessary for in vivo stability of the small subunit of the mammalian mitochondrial ribosome. Cell metabolism 9(4):386–397.

8. Seidel-Rogol BL, McCulloch V, & Shadel GS (2003) Human mitochondrial transcription factor B1 methylates ribosomal RNA at a conserved stem-loop. Nature genetics 33(1):23–24.

9. Metodiev MD, et al. (2014) NSUN4 is a dual function mitochondrial protein required for both methylation of 12S rRNA and coordination of mitoribosomal assembly. PLoS genetics 10(2):e1004110.

10. Spahr H, Habermann B, Gustafsson CM, Larsson NG, & Hallberg BM (2012) Structure of the human MTERF4-NSUN4 protein complex that regulates mitochondrial ribosome biogenesis. Proceedings of the National Academy of Sciences of the United States of America 109(38):15253–15258.

11. Dubin DT, Taylor RH, & Davenport LW (1978) Methylation status of 13S ribosomal RNA from hamster mitochondria: the presence of a novel riboside, N4-methylcytidine. Nucleic acids research 5(11):4385–4397.

12. Taylor RW & Turnbull DM (2005) Mitochondrial DNA mutations in human disease. Nature reviews. Genetics 6(5):389–402.

13. Sharoyko VV, et al. (2014) Loss of TFB1M results in mitochondrial dysfunction that leads to impaired insulin secretion and diabetes. Human molecular genetics 23(21):5733–5749.

14. Fernandez-Vizarra E, Berardinelli A, Valente L, Tiranti V, & Zeviani M (2007) Nonsense mutation in pseudouridylate synthase 1 (PUS1) in two brothers affected by myopathy, lactic acidosis and sideroblastic anaemia (MLASA). Journal of medical genetics 44(3):173–180.

15. Garone C, et al. (2017) Defective mitochondrial rRNA methyltransferase MRM2 causes MELAS-like clinical syndrome. Human molecular genetics 26(21):4257–4266.

16. Shinawi M, et al. (2011) 11p14.1 microdeletions associated with ADHD, autism, developmental delay, and obesity. American journal of medical genetics. Part A 155A(6):1272–1280.

17. Bradfield JP, et al. (2019) A Trans-ancestral Meta-Analysis of Genome-Wide Association Studies Reveals Loci Associated with Childhood Obesity. Human molecular genetics.

18. Schubert HL, Blumenthal RM, & Cheng X (2003) Many paths to methyltransfer: a chronicle of convergence. Trends in biochemical sciences 28(6):329–335.

19. Martin JL & McMillan FM (2002) SAM (dependent) I AM: the S-adenosylmethionine-dependent methyltransferase fold. Current opinion in structural biology 12(6):783–793.

20. Kimura S & Suzuki T (2010) Fine-tuning of the ribosomal decoding center by conserved methyl-modifications in the Escherichia coli 16S rRNA. Nucleic acids research 38(4):1341–1352.

21. Decatur WA & Fournier MJ (2002) rRNA modifications and ribosome function. Trends in biochemical sciences 27(7):344–351.

22. Fukasawa Y, et al. (2015) MitoFates: improved prediction of mitochondrial targeting sequences and their cleavage sites. Molecular & cellular proteomics: MCP 14(4):1113–1126.

23. Schaefer M, Pollex T, Hanna K, & Lyko F (2009) RNA cytosine methylation analysis by bisulfite sequencing. Nucleic acids research 37(2):e12.

24. Kozbial PZ & Mushegian AR (2005) Natural history of S-adenosylmethionine-binding proteins. BMC structural biology 5:19.

25. Haute LV, et al. (2019) METTL15 introduces N4-methylcytidine into human mitochondrial 12S rRNA and is required for mitoribosome biogenesis. Nucleic acids research.

26. Wei Y, et al. (2012) Crystal and solution structures of methyltransferase RsmH provide basis for methylation of C1402 in 16S rRNA. Journal of structural biology 179(1):29–40.

27. Sullivan LB, et al. (2015) Supporting Aspartate Biosynthesis Is an Essential Function of Respiration in Proliferating Cells. Cell 162(3):552–563.

28. Wang T, et al. (2015) Identification and characterization of essential genes in the human genome. Science 350(6264):1096–1101.

29. Klinge S & Woolford JL, Jr. (2019) Ribosome assembly coming into focus. Nature reviews. Molecular cell biology 20(2):116–131.

30. Popova AM & Williamson JR (2014) Quantitative analysis of rRNA modifications using stable isotope labeling and mass spectrometry. Journal of the American Chemical Society 136(5):2058–2069.

31. Shi Z, et al. (2019) Mettl17, a regulator of mitochondrial ribosomal RNA modifications, is required for the translation of mitochondrial coding genes. FASEB journal: official publication of the Federation of American Societies for Experimental Biology:fj201901331R.

32. Bournat JC & Brown CW (2010) Mitochondrial dysfunction in obesity. Current opinion in endocrinology, diabetes, and obesity 17(5):446–452.

33. Schulz TJ & Tseng YH (2013) Brown adipose tissue: development, metabolism and beyond. The Biochemical journal 453(2):167–178.

34. Shalem O, et al. (2014) Genome-scale CRISPR-Cas9 knockout screening in human cells. Science 343(6166):84–87.

35. Kehrein K, et al. (2015) Organization of Mitochondrial Gene Expression in Two Distinct Ribosome-Containing Assemblies. Cell reports 10(6):843–853.

36. Ma H, et al. (2019) N(6-)Methyladenosine methyltransferase ZCCHC4 mediates ribosomal RNA methylation. Nature chemical biology 15(1):88–94.

37. Rieder D, Amort T, Kugler E, Lusser A, & Trajanoski Z (2016) meRanTK: methylated RNA analysis ToolKit. Bioinformatics 32(5):782–785.

